# Deciphering Functional Redundancy in the Human Microbiome

**DOI:** 10.1101/176313

**Authors:** Liang Tian, Xu-Wen Wang, Ang-Kun Wu, Yuhang Fan, Jonathan Friedman, Amber Dahlin, Matthew K. Waldor, George M. Weinstock, Scott T. Weiss, Yang-Yu Liu

## Abstract

Although the taxonomic composition of the human microbiome varies tremendously across individuals, its gene composition or functional capacity is highly conserved^1-5^---implying an ecological property known as *functional redundancy*. Such functional redundancy is thought to underlie the stability and resilience of the human microbiome^6,7^, but its origin is elusive. Here, we investigate the basis for functional redundancy in the human microbiome by analyzing its genomic content network --- a bipartite graph that links microbes to the genes in their genomes. We show that this network exhibits several topological features, such as highly nested structure and fat-tailed gene degree distribution, which favor high functional redundancy. To explain the origins of these topological features, we develop a simple genome evolution model that explicitly considers selection pressure, and the processes of gene gain and loss, and horizontal gene transfer. We find that moderate selection pressure and high horizontal gene transfer rate are necessary to generate genomic content networks with both highly nested structure and fat-tailed gene degree distribution, and consequently favor high functional redundancy. These findings provide insights into the relationships between structure and function in complex microbial communities. This work elucidates the potential ecological and evolutionary processes that create and maintain functional redundancy in the human microbiome and contribute to its resilience.

---

The human microbiome harbors a plethora of taxa carrying distinct genes and gene families^4^, making it functionally diverse. At the same time however, the human microbiome is functionally redundant^7,8^, with many phylogenetically unrelated taxa carrying similar genes and performing similar functions^9-12^. For example, dietary carbohydrates can be metabolized by either *Prevotella* (from the phylum Bacteroidetes) or *Ruminococcus* (from the phylum Firmicutes)^13^. Short-chain fatty acids can be produced by multiple common genera including *Phascolarctobacterium, Roseburia, Bacteroides, Clostridium, Ruminococcus*, etc^14^. Bile acids can be modified by bacteria belonging to Lachospiraceae, Clostridiaceae, Erysipelotrichaceae and Ruminococcaceae^15^. Interleukin secretion can be promoted by *Sutterella, Akkermansia, Bifidobacterium, Roseburia* and *Faecalibacterium prausnitzii*^16,17^. Moreover, several metagenomic studies have reported that the carriage of microbial taxa varies tremendously within healthy populations, while microbiome gene compositions or functional profiles remain remarkably conserved across individuals^1-5^. Despite the functional variations and microbial gene diversity that have been uncovered through refined computational metagenomic processing^18^ and meta-analysis^19^, the highly conserved functional profiles across individuals imply significant functional redundancy (FR) in the human microbiome.

It has been suggested that this significant FR underlies the stability and resilience of the human microbiome in response to perturbations^6,7^, but there is little evidence to substantiate this idea. The origin of the FR observed in the human microbiome is still not well understood. A paradox has been raised recently, based on the fact that selection pressures could operate at different levels in the human-microbial hierarchy^20^, which potentially could drive the FR of the human microbiome in opposite directions. From the host perspective, although strong FR does not necessarily imply that the host is regulating the diversity of microbiota to promote FR^21^, host-driven or “top-down” selection would result in a community composed of widely divergent microbial lineages whose genomes contain functionally similar suites of genes, leading to high FR within the community. From the microbial perspective, however, competition between members of the microbiota would exert “bottom-up” selection pressure that results in specialized genomes with functionally distinct suites of genes, leading to high functional diversity (FD) and low FR within the community. This apparent paradox is oversimplified, as it doesn’t take into account the spatial structure and heterogeneous environments inhabited by the human microbiome. Nevertheless, low FR will tend to arise from widely divergent microbial lineages with functionally distinct suites of genes inhabiting the diverse niches within host body sites. On the other hand, high FR will arise from the presence of a core or common set of genes, i.e., housekeeping genes, required for diverse microbes to perform basic cellular functions and/or survive in the host body site they inhabit.

Here we investigate whether there is any organizing principle or assembly rule of the human microbiome that explains the observed high level of FR. In particular, we constructed the *genomic content network* (GCN) of the human microbiome, which enables us to quantify the within-sample FR for any given human microbiome sample for the first time. Then we applied tools from network science^22^ to study the topological features of the GCN that determine the FR of human microbiome samples. Furthermore, we developed a simple genome evolution model that can reproduce all the key topological features of the GCN. Using this model, we identified key evolutionary and ecological factors that account for the topological features of the GCN, and hence revealed the origin of FR in the human microbiome.

Consider a pool of *N* taxa, hereafter referred to as a metacommunity^23^, which contains a collection of *M* genes. The microbial composition or taxonomic profile 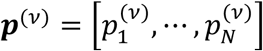. of a local community *ν* (i.e., a microbiome sample from a particular body site of a human subject *ν*) can be directly related to its gene composition or functional profile 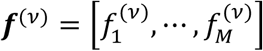. through the GCN of the metacommunity (Fig.1a-c). Here, we define the GCN as a weighted bipartite graph connecting these taxa to their genes. The GCN can be represented by an *N* × *M* incidence matrix ***G*** = (*G*_*ia*_), where a non-negative integer *G*_*ia*_ indicates the copy number of gene-*a* in the genome of taxon-*i* (Fig.1b). The functional profile is given by ***f***^(*v*)^ = *c****p***^(*v*)^ ***G***, where 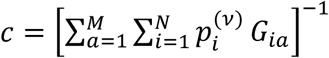. is a normalization constant.

**Figure 1:**
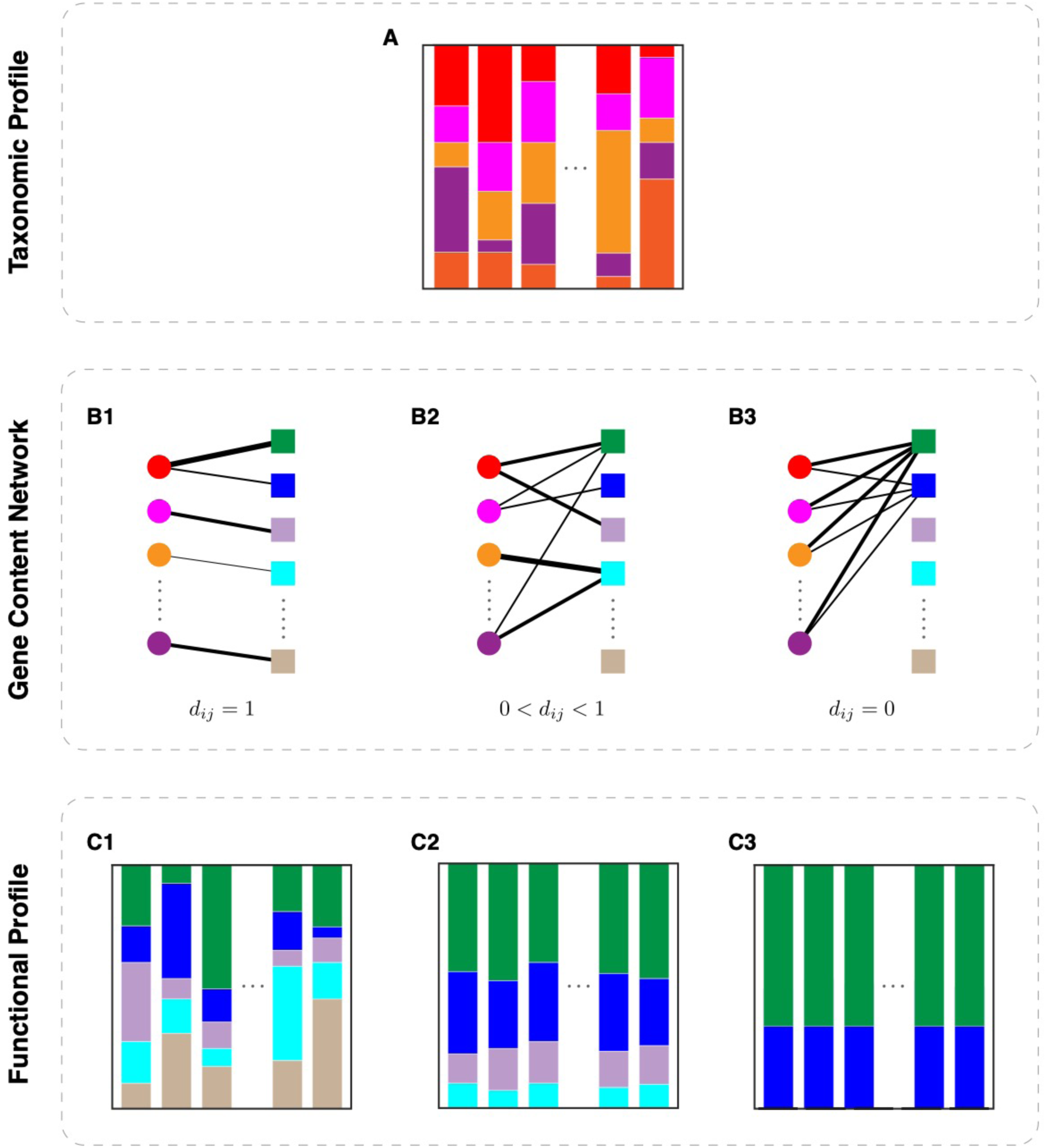
Structure of the genomic content network is crucial for determining the functional redundancy and functional diversity of microbial communities. Here we use hypothetical examples to demonstrate this point. **a**, The taxonomic profiles vary drastically across many local communities (i.e., microbiome samples from different individuals). **b**, Genomic content networks are bipartite graphs that connect taxa to the genes in their genomes. The left-hand side nodes (circles) represent different taxa and the right-hand side nodes (squares) represent different genes. The edge weight represents the gene copy number. **b-1**, Each taxon has a unique genome. **b-2**, Different taxa share a few common genes, some taxa are specialized to have some unique genes. **b-3**, All taxa share exactly the same genome. **c**, For each microbiome sample, its functional profile can be calculated from its taxonomic profile in (a) and the genomic content network in (b). **c-1**, The functional profiles vary drastically across different microbiome samples. For each sample, the functional diversity is maximized while the functional redundancy is minimized. **c-2**, The functional profiles are highly conserved across different samples. The within-sample functional diversity and functional redundancy are comparable. **c-3**, The functional profiles are exactly the same across all different microbiome samples. For each sample, the functional diversity is minimized while the functional redundancy is maximized.

A key advantage of GCN is that it enables us to calculate the FR for each local community, i.e., the within-sample or alpha functional redundancy (hereafter denoted as FR_*α*_). In the ecological literature, the FR of a local community is often interpreted as the part of its alpha taxonomic diversity (TD_*α*_) that cannot be explained by its alpha functional diversity (FD_*α*_) ^24-26^; i.e., FR_*α*_ ≡ TD_*α*_ − FD_*α*_. Typically, TD_*α*_ is chosen to be the Gini-Simpson index: 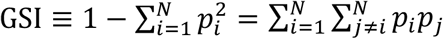, representing the probability that two randomly chosen members of the local community (with replacement) belong to two different taxa; and FD_*α*_ is chosen to be the Rao’s quadratic entropy 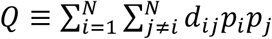, a classical alpha diversity measure that characterizes the mean functional distance between any two randomly chosen members in the local community^24,25^.

Here *d*_*ij*_ = *d*_*ji*_ ∈ [0, 1] denotes the functional distance between taxon-*i* and taxon-*j*, which can be calculated as the weighted Jaccard distance between the genomes of the two taxa. By definition, *d*_*ii*_ = 0 for *i* = 1, …, *N*. Note that with TD_*α*_ = GSI and FD_*α*_ = *Q*, we have 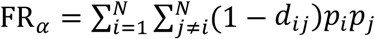, naturally representing the functional similarity (or overlap) of two randomly chosen members in the local community. Of course, one can also use other definitions for TD_*α*_ and FD_*α*_, then the expression of FR_*α*_ will be different. In particular, one can consider a parametric class of taxonomic (or functional) diversity measures based on Hill numbers^27,28^. We have confirmed that this does not affect our main results presented below.

The FR_*α*_ of each local community (or microbiome sample) is closely related to the system-level FR observed over a collection of samples. Consider two extreme cases: (i) Each taxon is completely specialized and has its own unique genome (Fig.1b1), hence *d*_*ij*_ = 1 for any *i ≠ j*. In this case, for each sample we have FD_*α*_ = TD_*α*_ and FR_*α*_ = 0. The functional profiles vary drastically across samples (Fig.1c1). (ii) All taxa share exactly the same genome (Fig.1b3), rendering *d*_*ij*_ = 0 for all *i* and *j*. In this case, for each sample we have FD_*α*_ = 0 and FR_*α*_ = TD_*α*_. The function profiles are exactly the same for all samples (Fig.1c3). These two extreme scenarios are of course unrealistic. In a more realistic intermediate scenario, the GCN has certain topological features such that different taxa share a few common functions, but some taxa are specialized to perform some unique functions (Fig.1b2). In this case, the FD_*α*_ and FR_*α*_ of each sample can both be high. Moreover, the functional profiles can be highly conserved across samples (Fig.1c2).

To quantitatively study the GCN underlying the human microbiome, we constructed a reference GCN using the Integrated Microbial Genomes & Microbiomes (IMG/M) database^29^, focusing on the Human Microbiome Project (HMP) generated metagenome datasets^30^. The IMG/M-HMP database used here includes in total 1,555 strains and 7,210 KEGG Orthologs (KOs). Here, each KO is a group of genes representing functional orthologs in molecular networks^31^. In order to reduce the culturing and sequencing bias for certain species (e.g., *Escherichia coli*), we randomly chose a representative strain (genome) for each species, which results in a reference GCN of 796 species and 7,105 KOs. This reference GCN is depicted in Fig.2a as a bipartite graph, where for visualization purposes each taxon node represents an order and each function node represents a KEGG super-pathway.

**Figure 2:**
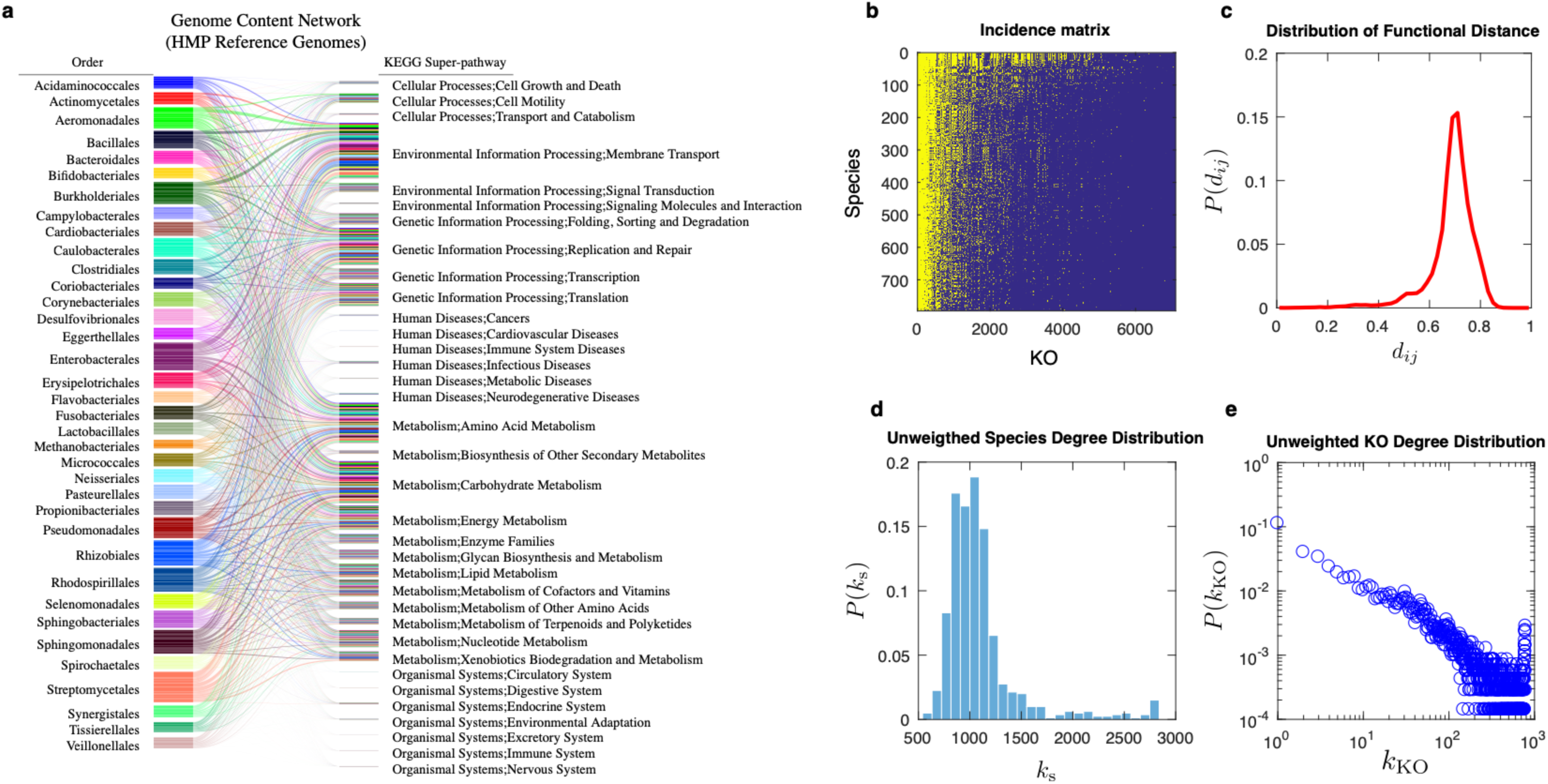
The genomic content network (GCN) constructed from the Integrated Microbial Genomes and Microbiome (IMG/M) database has nested structure and heterogeneous gene degree distribution. We use IMG/M-HMP, an IMG/M data mart that focuses on the Human Microbiome Project (HMP) generated metagenome datasets^30^ to construct the GCN. **a**, For visualization purpose, we depict this reference GCN at the order level for taxon nodes and at the KEGG super-pathway level for function nodes. The bar height of each order corresponds to the average genome size of those species belonging to that order. The thickness of a link connecting an order and a KEGG super-pathway is proportional to the number of KOs that belong to that super-pathway, as well as the genomes of species in that order. The majority of the super-pathways shown here are related to the metabolic, environmental, and genetic processes performed by microbes. However, for a small number of taxa, since some of their genes have mammalian and/or human disease orthologs, we also identified several super-pathways involved in human diseases and higher-order organizational systems. **b**, The incidence matrix of this reference GCN is shown at the species-KO level, where the presence (or absence) of a link between a species and a KO is colored in yellow (or blue), respectively. We organized this matrix using the Nestedness Temperature Calculator to emphasize its nested structure^32^. The nestedness value (∼0.34712) of this network is calculated based on the classical NODF measure^33^. **c**, The probability distribution of functional distances (*d*_*ij*_) among different species. The bin size is 0.02. **d**, The unweighted species degree distribution. Here, the unweighted degree of a species is the number of distinct KOs in its genome. e, The unweighted KO degree distribution. Here, the unweighted degree of a KO is the number of species whose genomes contain this KO.

In order to characterize the structure of this reference GCN, we systematically analyzed its network properties at the species-KO level. We first visualized its incidence matrix (Fig.2b), where the presence (or absence) of a link connecting a species and a KO is colored in yellow (or blue), respectively. We noticed that this matrix displays *a highly nested structure*^32-34^, i.e., the KOs of those species in the lower rows (with smaller genome size) tend to be subsets of KOs for those species in the higher rows (with larger genome size). The nestedness of the GCN can be quantified using the classical NODF measure^33^, and turns out to be much higher than expected by chance. We then calculated the functional distances among different species, finding a unimodal distribution with the peak centered around 0.7 (Fig.2c); finally, the unweighted degree distributions of taxon nodes (species) and function nodes (KOs) were calculated. Here, the unweighted degree of a species is just the number of distinct KOs in its genome, and the unweighted degree of a KO is the number of species whose genomes contain this KO. We found that the unweighted degrees of species follow a Poisson-like distribution (Fig.2d), implying that in general, species contain very similar numbers of distinct KOs. By contrast, the unweighted degree distribution of KOs is highly heterogeneous and displays a fat tail (Fig.2e), indicating that most KOs are specialized and only exist in the genomes of very few species, and a few housekeeping KOs appear in almost every species’ genome to maintain basic cellular functions. (Note that these housekeeping KOs also appear as the leftmost yellow columns in the incidence matrix shown in Fig.2b.) This is consistent with the characteristic asymmetrical U-shape observed in the gene frequency distributions of prokaryotic pangenomes^35,36^. Analyses of the reference GCN constructed by using other genome annotation, e.g., Clusters of Orthologous Groups of proteins (COGs)^37^, or constructed from a different database (MBGD: Microbial Genome Database for Comparative Analysis)^38^ revealed very similar network properties and did not affect our main results presented below.

The highly nested structure of the reference GCN is intriguing. This structure cannot be simply accounted for by housekeeping genes or the U-shape gene degree distribution. First, as shown in Fig.2b, the incidence matrix of the GCN still displays a highly nested structure even in the absence of housekeeping genes (the leftmost yellow columns). Second, if we randomize the GCN but preserve the gene degree distribution, the randomized GCNs have much lower nestedness than that of the real GCN. Third, we adopted tools from statistical physics to calculate the expected nestedness value and its standard deviation for an ensemble of randomized GCNs in which the expected species and gene degree distributions match those of the real GCN^39^. We found that the expected nestedness of randomized GCNs is significantly lower than that of the real GCN (one sample t-test yields *p*_value_ = 6.2853 × 10^−5^).

Using whole-genome shotgun (WGS) sequencing data from two large-scale microbiome studies, the Human Microbiome Project (HMP)^1,40,41^ and the MetaHIT (Metagenomics of the Human Intestinal Tract)^4,42^, we calculated the FR of human microbiome samples collected from different body sites. First, we constructed body site-specific GCNs using the IMG/M-HMP database. Note that the body site-specific GCNs display similar network properties as the global reference GCN constructed from the IMG/M-HMP database. To remove the potential impact of body site-dependent TD_*α*_ on the calculated FR_*α*_, we computed the normalized FR_*α*_ (i.e., nFR_*α*_ ≡ FR_*α*_*/*TD_*α*_) for these samples. Interestingly, we found that in both HMP and MetaHIT studies and for most body sites nFR_*α*_∼0.4 (Fig.3a-b, black boxes), suggesting that FR_*α*_ and FD_*α*_ are generally comparable for human microbiome samples. We also confirmed that the results are not sensitive to the integrity of the KEGG database, since nFR_*α*_ is stable if we randomly remove KOs from the GCN. Moreover, additional analyses demonstrated that although housekeeping KOs contribute to higher FR values, they are not the primary explanation for FR (Fig. S6).

**Figure 3:**
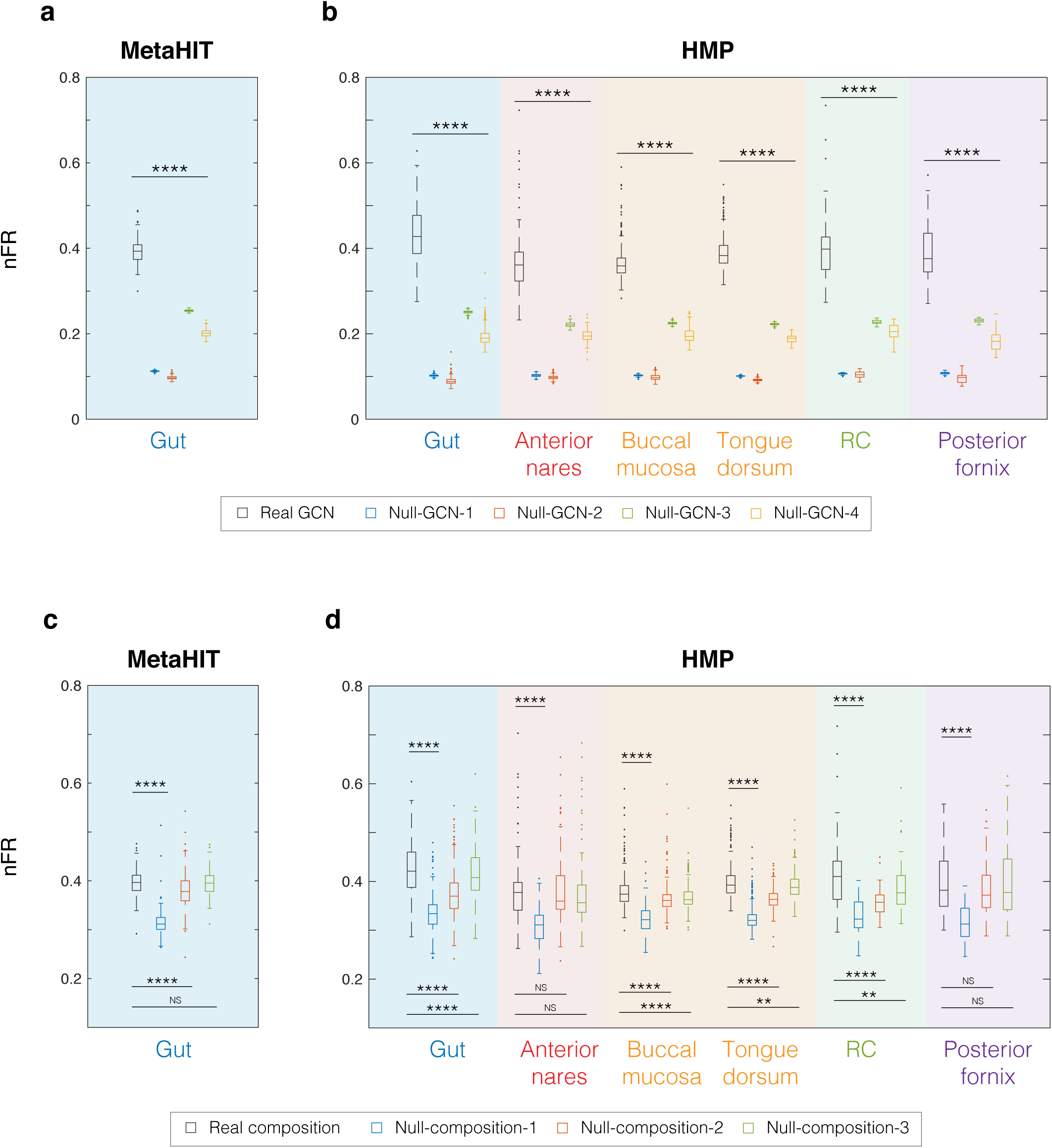
Topological features of the genomic content network and the assemblage pattern in the human-associated microbial communities contribute to the high functional redundancy observed in the human microbiome. Shotgun metagenomic data from HMP^1,40,41^ (for six different body sites) and MetaHIT^4,42^ (for gut) were analyzed. **a-b**, The box plots of the normalized function redundancy (nFR ≡ FR/TD) were calculated from the real GCN (black box), as well as the randomized GCNs (colored boxes) using four different randomization schemes: Complete randomization (Null-GCN-1); Species-degree preserving randomization (Null-GCN-2); KO-degree preserving randomization (Null-GCN-3); Species- and KO-degree preserving randomization (Null-GCN-4). Here the (weighted) degree of a KO is the sum of copy numbers of this KO in those genomes that contain it, and the (weighted) degree of a species is the sum of copy numbers of those KOs in this species’ genome. **c-d**, The box plots of normalized function redundancy were calculated from the real microbial compositions (black box), as well as the randomized microbial compositions (colored boxes) using three different randomization schemes: Randomized microbial assemblage generated by randomly choosing the same number of species from the species pool but keeping the species abundance profile unchanged (Null-composition-1); Randomized microbial abundance profiles through random permutation of non-zero abundance for each sample across different species (Null-composition-2); Randomized microbial abundance profiles through random permutation of non-zero abundance for each species across different samples (Null-composition-3). Significance levels: FDR<0.05 (*), <0.01(**), <0.001(***), <0.0001(****); >0.05 (NS, non-significant); Wilcoxon signed-rank test.

To identify key topological features of the GCN that determine nFR_*α*_, we adopted tools from network science. In particular, we randomized the body site-specific GCNs using four different randomization schemes, yielding four different null models. Then, we recalculated nFR_*α*_ for each sample (Fig.3a-b, colored boxes), finding that for all the body sites examined all the four different null models yield lower nFR_*α*_ than those calculated from real body site-specific GCNs (Fig.3a-b, black boxes). Analyzing the network properties of those null models (Figs.S7-S8), we found that those randomized GCNs all display lower nestedness and higher ⟨*d*_*ij*_⟩ than those of the real GCNs. Thus, the highly-nested structure and low ⟨*d*_*ij*_⟩ of the real GCNs contribute to the high nFR_*α*_ values observed in the microbiome samples. Moreover, for the first two null models (Null-GCN-1 and Null-GCN-2, where both the highly-nested structure and high gene degree heterogeneity of the real GCN are destroyed), nFR_*α*_ is much lower than those of the other two null models (Null-GCN-3 and Null-GCN-4, where the highly-nested structure is destroyed, but the high gene degree heterogeneity is kept). This suggests that the high gene degree heterogeneity also contribute to the high nFR_*α*_ values of those microbiome samples. Hence the GCN exhibits at least three different topological features (highly nested structure, low ⟨*d*_*ij*_⟩, and heterogeneous gene degree distribution) that jointly contribute to the high nFR_*α*_ value of microbiome samples. We emphasize that these findings do not depend on the detailed definitions of *d*_*ij*_, FR_*α*_, FD_*α*_, or the functional annotation of genomes (Figs.S1, S2, S4).

To test if the microbe assemblages or their abundances play an important role in determining nFR_*α*_, we randomized the taxonomic profiles using three different randomization schemes, yielding three different null models. Then, we recalculated nFR_*α*_ for each sample (Fig.3c-d, colored boxes). We found that for each microbiome sample if we preserve the abundance profile but randomly replace the species by those present in the species pool (i.e., in the corresponding body site-specific GCN), the resulting null composition model (Null-compostion-1) always yields much lower nFR_*α*_ than that of the original sample. This suggests that the species present in each microbiome sample or local community are not assembled at random, but follow certain functional assembly rules^44^. Interestingly, if we randomize the microbial compositions through random permutation of non-zero abundance for each sample across different species (Null-composition-2) or for each species across different samples (Null-composition-3), those two null models did not always yield much lower nFR_*α*_ than that of the original sample. Again, these observations do not rely on the detailed definitions of *d*_*ij*_, FR_*α*_, FD_*α*_, or the functional annotation of genomes (Figs.S1, S2, S4). These observations suggest that the assemblage of microbes plays a more important role than their abundances in determining the high FR of the human microbiome. We hypothesize that the specific environment (e.g., the host nutrient and immune state) from particular microbiome samples were obtain will tend to select for sets of functions among most or all inhabitants, at any abundance. This could partially explain why assemblage or membership matters more than abundances in determining functional redundancy.

All the results calculated from WGS data presented above are based on taxonomic profiling using existing reference genomes. To test if our findings could be derived independent of reference genomes, we adopted a *de novo* method to perform taxonomic profiling of shotgun metagenomic data without using any reference genomes^42^. This *de novo* taxonomic profiling method is based on the binning of co-abundant genes across a series of metagenomic samples. We applied this method to the human gut microbiome samples from MetaHIT to construct a GCN. Notably, we found that this GCN displays very similar network properties as the GCN constructed using reference genomes, i.e., high nestedness, a unimodal functional distance distribution with a clear peak centered around 0.7, Poisson-like species degree distribution, and a fat-tailed gene degree distribution. Using the taxonomic profiles and the constructed GCN obtained from this method, we further calculated the normalized functional redundancy of real microbiome samples and compared these values to those calculated from randomized GCNs or randomized microbial compositions. We found that all the key findings presented in Fig.3 can be reproduced, implying that our results do not depend on the existing reference genomes.

To gain more biological insight into the bases of the topological features of the real GCN, and thus deepen understanding of the origin of FR in the human microbiome, we developed a simple genome evolution model. In this model, we explicitly considered selection pressure and the processes of gene gain and loss, and horizontal gene transfer (HGT) (Fig.4a). We assumed selection pressure simply favors changes in larger genomes. We found that with reasonable model parameters all the key topological features of the real GCN can be reproduced by our simple model (Fig.4b-e). Moreover, we found that a high HGT rate is necessary to generate a GCN with a highly nested structure (Fig.4f) and a very heterogeneous gene degree distribution as observed in the real GCN (Fig.4g), which are crucial features to maintain high FR in the human microbiome. As shown in Fig.4f, the nestedness (measured by NODF) of the GCN generated by our model displays a phase-transition like behavior: when the HGT rate is above certain threshold value, NODF deviates from zero and increases gradually. Similarly, as shown in Fig.4g, the Kullback–Leibler (KL) divergence between the normalized gene degree distribution of real GCN and that of a simulated GCN also displays a phase-transition like behavior. When the HGT rate is above certain threshold value, the KL divergence drops and becomes very close to zero, implying that the gene degree distribution of the generated GCN is very similar to that of the real GCN. These results highlight the importance of HGT in determining the high FR of the human microbiome. We further demonstrated that both the incidence matrix of GCN and the functional distance distribution will be quite different from that observed in the real GCN, if the selection pressure is zero or too large. This implies that moderate selection pressure is needed to reproduce key topological features of the GCN, and consequently favors high functional redundancy.

**Figure 4:**
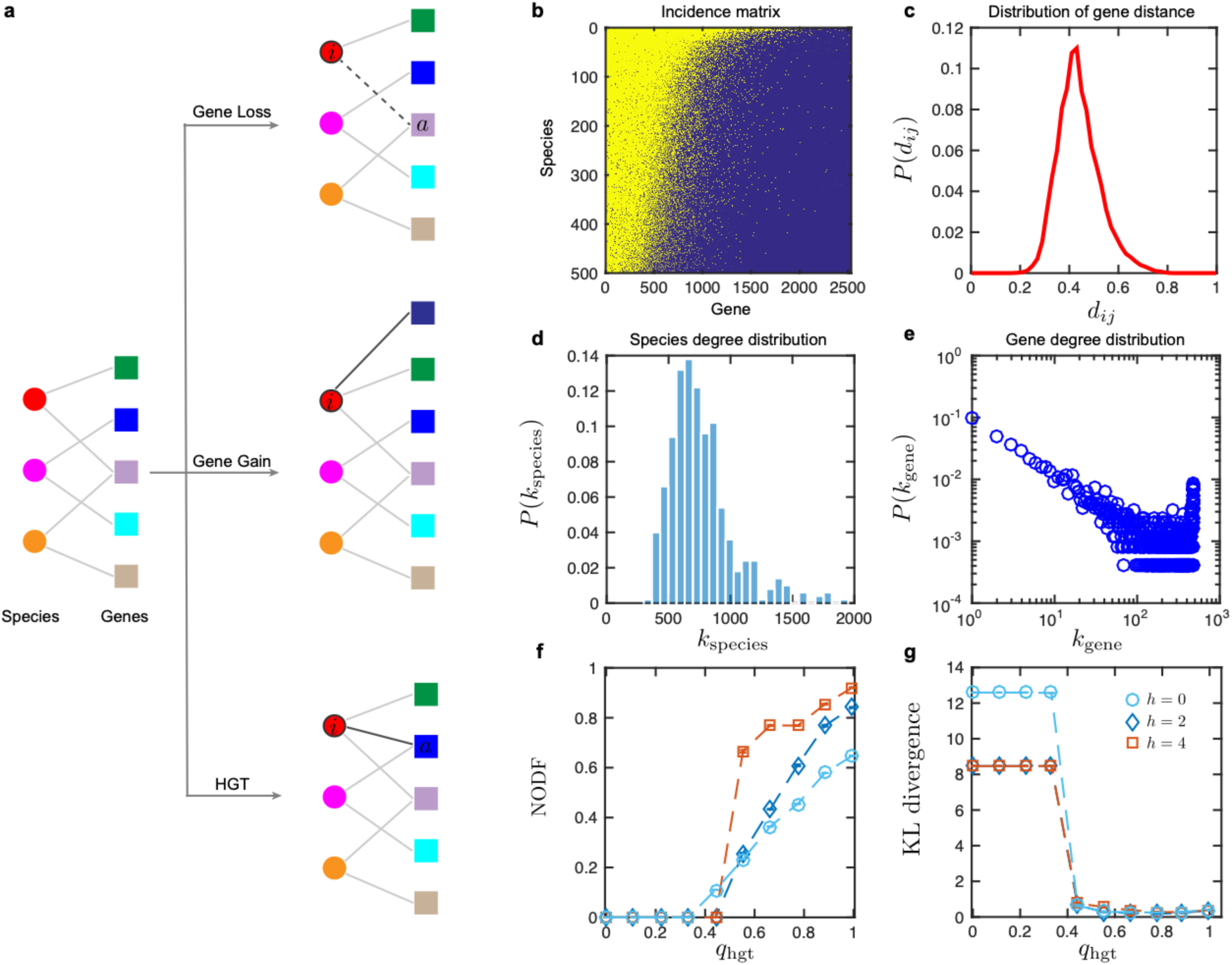
A simple genome evolution model can generate GCNs that capture key topological features of the real GCN. **a**, Schematic diagram of the genome evolution model. At each time step *t*, the genome of a species *i* (shown in red) randomly chosen with probability proportional to 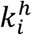 will be updated based on one of the following three events: gene loss, gene gain, and horizontal gene transfer (HGT), with corresponding rates *q*_gl_, *q*_gg_, *q*_HGT_, respectively. Note that the parameter *h* ≥ 0 representing the selection pressure, and *h* = 0 corresponds to the case of neutral model. The three rates naturally satisfy *q*_gl_ + *q*_gg_ + *q*_HGT_ = 1. During HGT, a gene *a* from a randomly chosen donor species is randomly selected and then transferred to the genome of species *i*. During gene loss, a gene *a* in the genome of species *i* is randomly selected and then removed. During gene gain, a new gene is added to the genome of species *i*. The initial GCN is a random bipartite graph that consists of 500 species and 200 genes with connection probability 0.8. The total number of evolution time steps is 5×10^5^. **b-e**, The incident matrix of the final GCN (with nestedness value NODF=0.703), functional distance, species degree and gene degree distributions with *h* = 2, *q*_HGT_ = 0.795, *q*_gg_ = 0.005, and *q*_gl_ = 0.2. **f**, The nestedness (quantified by NODF) of the final GCN calculated as a function of HGT rate with different selection pressure *h* = 0, 2, 4. **g**, The Kullback–Leibler (KL) divergence between the normalized gene degree distribution 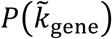 of real GCN and that of the simulated GCNs calculated with different selection pressures and HGT rates as shown in panel-f. Here the normalized gene degree 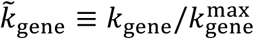.

In sum, we developed a GCN based framework to quantify the FR of the human microbiome and revealed the origin of FR using a genome evolution model. The GCN framework will enable us to directly validate if a strong FR underlies the stability and resilience of the human microbiome in response to perturbations^6,7^, such as probiotic administration and fecal microbiota transplantation. This could potentially inform microbiome-based therapies, if FR can indeed serve as a residence indicator of the human microbiome. FR has been found in many other microbial systems as well, e.g., plant microbiome^45,46^, ocean microbiome^47^, and soil microbiome^48,49^. Our general, quantitative measure of FR can also be directly applied to those microbial systems and hence facilitate a direct test of the hypothesis that there are systematic differences in FR between free-living and host-associated microbial communities^50^. More broadly, we anticipate that the GCN framework will yield new insights into the relationships between biodiversity and ecosystem function for diverse microbial communities.

## Acknowledgements

We thank Yandong Xiao, Eugene Koonin, Patrick Chain, Uri Gophna for valuable discussions. Special thanks to Brigid Davis and Edwin K. Silverman for a careful reading of the manuscript.

## Author Contributions

Y.-Y.L conceived and designed the project. L.T., X.-W.W., A.-K.W., Y.-H.F. and Y.-Y.L. did the analytical and numerical calculations. L.T. analyzed all the empirical data. X.-W.W. developed the genome evolution model. All authors interpreted the results. Y.-Y.L. wrote the manuscript. L.T., X.-W.W., A.-K.W., Y.-H.F., J.F., M.K.W and S.T.W. edited the manuscript.

## Author Information

The authors declare no competing financial interests. Correspondence and requests for materials should be addressed to Y.-Y.L.

